# Resting-state and task-based centrality of dorsolateral prefrontal cortex predict resilience to inhibitory repetitive transcranial magnetic stimulation

**DOI:** 10.1101/735506

**Authors:** Sophie M.D.D. Fitzsimmons, Linda Douw, Odile A. van den Heuvel, Ysbrand D. van der Werf, Chris Vriend

**Author notes:** Corresponding author details: Name: Sophie M.D.D. Fitzsimmons, Postal address: Human and Life Sciences Building 13W55, De Boelelaan 1108, 1081 HZ Amsterdam, The Netherlands.

## Abstract

**Background:** Repetitive transcranial magnetic stimulation (rTMS) is used to investigate normal brain function in healthy participants and as a treatment for brain disorders. Various subject factors can influence individual response to rTMS, including brain network properties.

**Objective/Hypothesis:** A previous study by our group showed that ‘virtually lesioning’ the left dorsolateral prefrontal cortex (dlPFC; important for cognitive flexibility) using inhibitory rTMS reduced performance on a set-shifting task. We aimed to determine whether this behavioural response was related to topological features of pre-TMS resting-state and task-based functional networks.

**Methods:** Inhibitory (1Hz) rTMS was applied to the left dlPFC in 16 healthy participants, and to the vertex in 17 participants as a control condition. Participants performed a set-shifting task during fMRI at baseline and directly after a single rTMS session 1-2 weeks later. Functional network topology measures were calculated from resting-state and task-based fMRI scans using graph theoretical analysis.

**Results:** The dlPFC-stimulated group, but not the vertex group, showed reduced set shifting performance after rTMS associated with lower task-based betweenness centrality of the dlPFC at baseline (p=.030) and a smaller reduction in task-based betweenness centrality after rTMS (p=.024). Reduced repeat trial accuracy after rTMS was associated with higher baseline resting state node strength of the dlPFC (p=.017).

**Conclusions:** Our results suggest that behavioural response to inhibitory rTMS to the dlPFC is dependent on baseline functional network features. Individuals with more globally integrated stimulated regions show greater resilience to inhibitory rTMS, while individuals with more locally well-connected regions show greater vulnerability.

**Highlights:**

- Functional brain network properties predict behavioural response to 1Hz DLPFC rTMS
- Globally integrated stimulated regions are resilient to effects of inhibitory rTMS
- Segregated, locally well-connected regions are vulnerable to inhibitory rTMS
- Change in performance after rTMS correlates with change in network properties

## Introduction

Repetitive transcranial magnetic stimulation (rTMS) is a method of non-invasively exciting (using high frequency (HF) stimulation, >5Hz) or inhibiting (using low frequency (LF) stimulation, ≤1 Hz) specific brain regions and connected networks through electromagnetic induction. It is used to investigate brain function in healthy subjects [1] and is becoming a common treatment in neurological and psychiatric patient populations[2]. However, individuals vary considerably in their response to rTMS. This variation in response is associated with a number of factors, including baseline structural[3] and functional connectivity[4] (FC) of the stimulated brain network.

FC is a measure of the temporal correlation of activity between anatomically separate brain areas[5]. FC of the targeted area is predictive of the outcome of rTMS to the dorsolateral prefrontal cortex (dlPFC) for the treatment of depression[4,6,7], dorsomedial PFC for the treatment of eating disorders[8], and of change in motor-evoked potential amplitude after rTMS of the motor cortex[9]. However, given that rTMS also influences the activity of areas distant to the stimulated site[10], metrics that take the organisation of the wider functional network into account may be a more useful predictor of rTMS outcome than seed-based or region of interest (ROI)-based FC, which are limited to measuring FC between *a priori*-defined brain regions.

An alternative method is to define the brain as a network consisting of nodes (brain regions) and edges (functional connections between regions), and then apply graph theoretical analysis to evaluate the organisation and topology of this network [11]. Graph measures can be extracted from the whole network, individual nodes, or subnetworks of more densely interconnected regions (modules[12]), allowing characteristics of the network to be evaluated at different spatial scales. While functional network features have been shown to be predictive of HF rTMS outcome in depression[13], this method has not yet been applied to the prediction of cognitive outcomes of LF rTMS.

In an earlier study by our group in healthy participants [14], we applied LF rTMS to the left dlPFC, a region important for executive function, specifically the ability to flexibly adapt to changes in rules or environment[15,16]., Following rTMS, subjects showed a reduction in performance on a set-shifting task (testing cognitive flexibility), during fMRI. In the present exploratory re-analysis of resting-state and task-based fMRI data acquired before and after rTMS, we aimed to determine whether baseline functional network characteristics are predictive of behavioural response to inhibitory rTMS, and whether TMS-induced change in network characteristics is associated with change in performance.

We limited our choice of graph measures to those known to be markers of resilience and vulnerability to lesions, as we were investigating LF rTMS that causes a temporary ‘virtual lesion’. *Centrality* (i.e. how well-connected a node is[12]) is an important determinant of network resilience[17]. We applied three different centrality measures to the stimulated region: node strength (NS) that describes the total strength of a node’s connections, and is considered to be an indicator of local connectivity; betweenness centrality (BC), that measures how many high strength paths in the network pass through a node; and participation coefficient (PC), which describes whether a node is connected mostly to its own module or to other modules[12]. Different types of centrality may have different implications for the rTMS-induced behavioural effects. Since the loss of highly locally connected nodes (such as those with high NS) is predictive of network disruption[17,18], we predicted that this measure may be associated with greater vulnerability to the effects of inhibitory rTMS. On the other hand, nodes with high global connectivity (such as those with high BC or PC) are associated with greater cognitive flexibility[19], suggesting that they may be resilient to cognitive disruption. Therefore, we expected that higher BC and PC of the stimulated node would be associated with higher resilience to inhibitory rTMS.

## Material & Methods

### Participants

Our sample consisted of 33 healthy participants originally recruited for a previous study[20]. Participants were not included if they suffered from neurological or psychiatric illnesses, substance abuse, cognitive deficits, or had a family history of epilepsy. Sixteen participants (mean age of 55 ± 9 years, 9 men) were randomly appointed to dlPFC (verum) rTMS and 17 age and gender matched participants (mean age of 57 ± 10 years, 11 men) to vertex (active control) rTMS. All participants were screened for the presence of psychiatric disorders using the Structured Clinical Interview for DSM-IV Axis-I Disorders[21], depressive symptoms using the Beck Depression Inventory[22], anxiety symptoms using the Beck Anxiety Inventory[23], and general cognitive status using the Mini-Mental State Examination (MMSE)[24]. We used the Dutch version of the national adult reading test[25] to provide an estimate of intelligence. The study protocol was reviewed and approved by the Research Ethics Committee of the VU University medical center (VUmc) and all participants provided written informed consent.

### Experimental procedure

Full details of the experimental procedure and set-shifting task are reported in Gerrits et al 2015[14]. In brief, participants performed a set-shifting task during two fMRI sessions separated by an average of 16.9 ± 11.2 and 15.9 ± 7.7 days in the verum and control condition, respectively (figure 1A). Participants received inhibitory rTMS to either the dlPFC or vertex directly prior to the second fMRI session. For participants in the verum group, fMRI data acquired during a set-shifting task in this first session were used to determine coordinates for TMS coil localisation using neuro-navigation software (ASA4.1 software, ANT Neuro, The Netherlands). Specifically, the peak voxel of the switch>repeat contrast was used – group mean MNI coordinates were x=-42, y=28, z=31 (see below for more details about the set shifting task). For participants in the control condition, we used individual anatomical T1-weighted MR scans to determine the location of the vertex (mean stimulated MNI coordinates x=0, y=-34, z=70). During the second session, 1Hz rTMS was applied for 20 minutes (1200 pulses total) using a hand-held figure-of-eight TMS coil (Medtronic MagOption, Medtronic Denmark A/S, Copenhagen, Denmark) at 110% of the individual motor threshold to ‘virtually lesion’ the dlPFC or vertex. Participants then performed the set-shifting task for a second time during fMRI. The median interval between stimulation and the beginning of the set-shifting task was 5’19’’ for the verum and 5’56’’ for the control condition.

**Figure 1:**
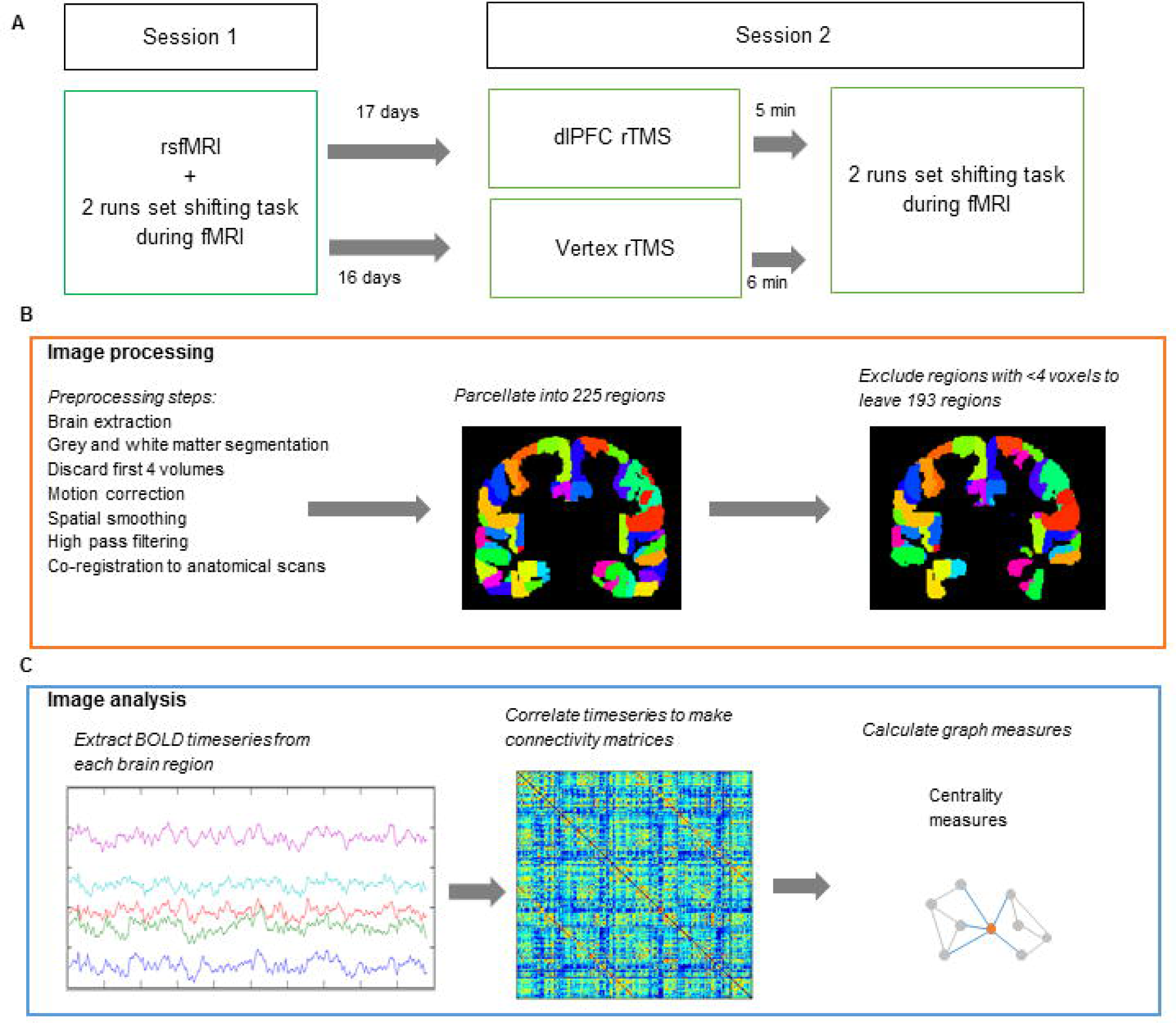
Study design and image processing/analysis. **A.** *Study design*. All participants attended two sessions. During the first session, participants underwent an rsfMRI scan and carried out a set-shifting task during two separate runs of fMRI. Participants were then randomised to either dlPFC or vertex rTMS groups. 16-17 days later, participants attended the second session. They received either dlPFC or vertex rTMS, followed directly by carrying out the set shifting task during 2 separate runs of fMRI. **B**. *Image processing steps*: fMRI scans were preprocessed and parcellated into 225 regions. This was followed by exclusion of regions containing <4 voxels. **C.** *Image analysis steps:* FC and graph theoretical analysis: The BOLD timeseries was extracted from each parcellated region. Pearson correlations were carried out between each pair of regions, giving a 193×193 correlation matrix for each participant. These matrices were then used to calculate centrality graph measures.

### Set-shifting task and behavioural data

During the set-shifting task (programmed in E-Prime version 2.0, Psychology Software Tools, Sharpsburg, PA, USA), an arrow appeared either on one of four sides of a fixation cross in the centre of the screen, pointing either down, up, left or right. The participant had to respond by pressing the up, down, left or right key depending on whether the current classification rule for the arrow was *location* or *direction*. The participant was informed about incorrect repeat trials or a change in the classification rule (i.e. set-shift) by the presentation of a red screen. A green screen signalled a correct response. The task continued until the participant had completed 40 correct set shift trials. Two versions of the set-shifting task were counterbalanced between the first and second session that differed in the starting location and orientation of the arrow. All behavioural responses were recorded using an MRI compatible response box. The task was practiced prior to data collection to obtain a stable level of performance. Each response during the task was classified as correct repeat, incorrect repeat, successful shift, incorrect shift, or no shift/no repeat. Data from these responses (reaction time (RT) and error rate (ER)) were used to calculate the following behavioural outcome measures:

- Percentage change in repeat reaction time (RRT) and shift reaction time (SRT) from baseline:

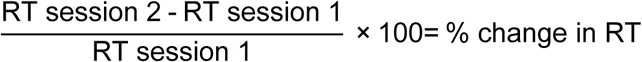
- Percentage change in repeat error rate (RER) and shift error rate (SER) from baseline. A constant (*c*) was added to the ER of session 1 when calculating percentage change to prevent dividing by zero:

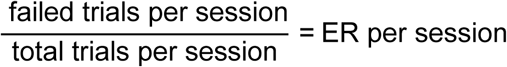

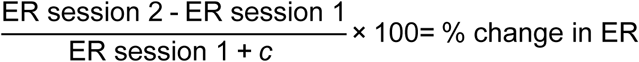

### Image acquisition

Functional imaging was performed at the VUmc, Amsterdam using a GE Signa HDxt 3-T MRI scanner (General Electric, Milwaukee, USA) using whole-brain gradient echo-planar imaging (EPI) sequences. Eyes-closed rsfMRI images (TR 1,800ms; TE = 30ms; 64×64 matrix, flip angle = 80°), were acquired with 40 ascending slices per volume (3.75 × 3.75 mm in-plane resolution; slice thickness = 2.8 mm; inter-slice gap = 0.2 mm). At baseline, rsfMRI scans were acquired before task-based scans. Functional images during the set-shifting task (TR=2,100ms; TE=30ms; 64×64 matrix, flip angle=80°) with 40 ascending slices per volume (same resolution as rsfMRI) were acquired in two runs; runs varied in length between participants, as the set shifting task lasted for as long as it took to achieve 40 correct set shifting trials. In the post-TMS task-based fMRI, the dlPFC group took on average 9.13 ± 0.95 mins and 8.17 ± 0.39 mins for the first and second run, respectively. The control group 9.17 ± 0.70 and 8.16 ± 0.61 mins. A sagittal 3D gradient-echo T1-weighted sequence (256×256 matrix; voxel size=1×0.977×0.977mm; 172 sections) was also acquired for co-registration and parcellation.

### fMRI analysis

We chose to analyse the task-based scans as two separate runs in order to carry out a replication of our own analyses. Image processing of the two runs of the task scan and rsfMRI scan was performed using FMRIB’s software library version 5.0.8 (FSL)[26] (figure 1B) and included discarding the first four volumes of the functional scan to reach magnetization equilibrium, motion correlation, 5 mm spatial smoothing, and high-pass filtering (see Appendix A for more details). The brain was parcellated into 225 brain regions using 210 cortical regions from the Brainnetome atlas[27], 14 individually segmented subcortical areas and one cerebellar ROI from FSL’s cerebellar atlas [28]. To account for EPI distortions near air/tissue boundaries during scanning, we excluded any nodes with less than four signal containing voxels [29]. A total of 193 regions common to all fMRI runs remained: excluded regions were located in the orbitofrontal gyrus, inferior temporal gyrus, parahippocampal gyrus, and thalamus.

### Graph analyses

Connectivity analyses (figure 1C) were performed using in-house scripts and the Brain Connectivity Toolbox[12] in MATLAB R2012a (The MathWorks, Inc, Natick, MA, USA). A connectivity matrix was created for each subject by calculating Pearson correlation coefficients between time-series from all nodes. All weights in the connectivity matrix were absolutized. Using the absolute values prevents loss of important interactions between brain regions[30] and using weighted rather than thresholded or binarised connectivity matrices, prevents discarding weaker but potentially relevant connections[31]. We carried out additional motion correction by ‘scrubbing’ all time points with >0.5 mm of framewise displacement prior to making the correlation matrices (as recommended by Power et al[32]).

Graph measures were calculated on a global and nodal scale. In order to identify the dlPFC node for each subject, we defined spherical ROIs (5 mm radius) corresponding to the coordinates of the stimulated brain area, and selected the atlas region that the ROI overlapped with most. A node in the visual cortex (node 202 from the Brainnetome Atlas, left V5) was also selected as a control node. We hypothesised that this node was unlikely to be affected by dlPFC rTMS, and was the node that appeared least frequently across individual participants’ dlPFC-containing modules as calculated by Louvain modularity (see Appendix B for more details).

We calculated the following centrality graph measures in both groups for the dlPFC node and the control node in the visual cortex [12]:

- Betweenness centrality (BC): the fraction of strongest weighted shortest paths that pass through a given node. This suggests that high BC nodes will be well connected throughout the entire network[12].
- Participation coefficient (PC): an assessment of the type of connections a node has. Low PC indicates more high weighted connections with one’s own module; high PC more high weighted connections to other modules[33].
- Node strength (NS): the sum of all edge weights, indicating how strongly connected a node is to its neighbours[12].

### Statistical analyses

Statistical analyses were carried out in SPSS Statistics 22 (IBM Corp., Armonk, NY, USA) and R. Independent samples t-tests or Mann-Whitney U tests (two tailed, α=0.05) were used to compare demographic and behavioural characteristics of verum and control condition, depending on the distribution. Since graph measures were not normally distributed, nonparametric tests were used for statistical analysis. Mann-Whitney U tests were used to compare baseline and rTMS-induced changes in graph measures between the verum and control group. Correlations between graph measures and behavioural outcomes were carried out using Kendall’s Tau-b correlations with bootstrapped 95% confidence intervals (CIs). Due to high levels of correlation between network measures and the exploratory nature of this study, these analyses were not corrected for multiple comparisons. The data did not meet the assumptions for regression analysis; therefore, to compare Kendall’s tau coefficient between verum and control groups we first converted Kendall’s tau to Pearson’s *r* using the formula r = sin (0.5 π τ)[34] and calculated z scores using a Fisher’s *r*- to-Z transform[35].

## Results

### Demographic characteristics and session information

The verum and control groups were well matched in terms of age (p=.606), sex (p=.619), and MMSE (p=.360), but the vertex group had a higher estimated intelligence (p=.012) (Table 1). The interval between session 1 and session 2 (p=.930) and between rTMS and task (p=.053) did not differ significantly between groups (Table 1).

**Table 1:** Demographic characteristics, session information and change in set shifting performance after rTMS

|  | *dIPFC group*<br>*n=16* | *Vertex group*<br>*n=17* | *p* |
| --- | --- | --- | --- |
| ***Demographics*** |  |  |  |
| Age (years) | 55 ± 9 (39-75) | 57 ± 10 (41-70) | .606<sup>a</sup> |
| Sex (no./% men) | 9/56% | 11/65% | .619<sup>b</sup> |
| Level of education reached <sup>c</sup><br>(median, range, % > 5) | 6 (4-7) 69% | 6 (3-7) 59% | .721<sup>b</sup> |
| Estimated IQ | 98 ± 12 (73-123) | 110 ± 14 (82-130) | **.012** |
| MMSE | 29 ± 1 (28-30) | 29 ± 1 (27-30) | .360 |
| ***Session information*** |  |  |  |
| Interval session 1 - session 2<br>(days) | 17 ± 11 (7-56) | 16 ± 8 (7-35) | .772<sup>a</sup> |
| Interval rTMS - task (s) | 319 ± 54 (240-488) | 356 ± 137 (277-800) | .053 |
| ***Change in set shifting performance after rTMS (% change from baseline)*** |  |  |  |
| RRT | -2.44 ± 14.70 (-30.65-33.41) | -6.00 ± 17.72 (-30.47-27.40) | .488 |
| SRT | -5.08 ± 8.01 (-21.75-14.36) | -9.17 ± 15.59 (-38.93-18.95) | .465 |
| RER | -0.26 ± 0.54 (-1.50-0.73) | -0.06 ± 0.65 (-1.09-1.67) | .533 |
| SER | 0.43 ± 0.96 (-1.13-3.12) | -0.27 ± 1.05 (-2.56-1.37) | **.049** |

Values are presented as mean ± standard deviation (range) unless otherwise indicated
Significance of group differences tested using an Independent Samples Mann-Whitney U test unless otherwise indicated.
<sup>a</sup> Independent samples t-test
<sup>b</sup> Pearson's $\chi^2$ test.
<sup>c</sup> Level of education expressed as the Verhage 7-point scale[47]: 1=no finished education; 5=secondary school, medium level; 7=university training. Proportions of people scoring ≤ 5 and >5 compared using $\chi^2$ test.
dIPFC, dorsolateral prefrontal cortex; MMSE, mini-mental state examination; RRT, repeat response time; SRT, switch response time; RER, repeat error rate; SER, switch error rate.

### Change in set shifting performance after rTMS

Groups did not differ in set shifting performance at baseline[14] (see also Appendix C, Table S1). There was a small rTMS-induced increase in shift trial error rate (SER) in the verum group compared with the control group (p=.049, Table 1). There were no other significant between-group differences in rTMS induced changes in set shifting performance. For both groups there was, in general, an improvement in performance between session 1 and session 2 in all behavioural measures except for SER. None of the changes in behavioural outcomes correlated with intelligence, educational level, MMSE or rTMS-task interval (see Appendix C, table S2).

### Prediction of behaviour change after rTMS from baseline resting state graph measures

The verum and control groups did not differ significantly in any resting state graph measures at baseline (see Appendix C, table S3). In the verum group, there was a positive correlation between the change in repeat error rate (ΔRER) and baseline NS of the dlPFC node (τ= 0.447, p=.017, 95% CI [0.138, 0.781]) (figure 2A) – i.e. the higher the pre-stimulation NS, the greater the increase in repeat errors after rTMS. This association was not seen in the control group (τ= −0.170, p=.343, 95% CI [−0.527, 0.214]), and the association in the verum group was also significantly stronger than that seen in the control group (z=2.696, p=0.007). In the verum group there was no significant association between NS in the control node within the visual cortex and ΔRER (τ=0.380, 95% CI [−0.049, 0.698]). None of the other resting state graph measures examined showed significant correlations with change in cognitive performance (see Appendix C, table S4).

**Figure 2:**
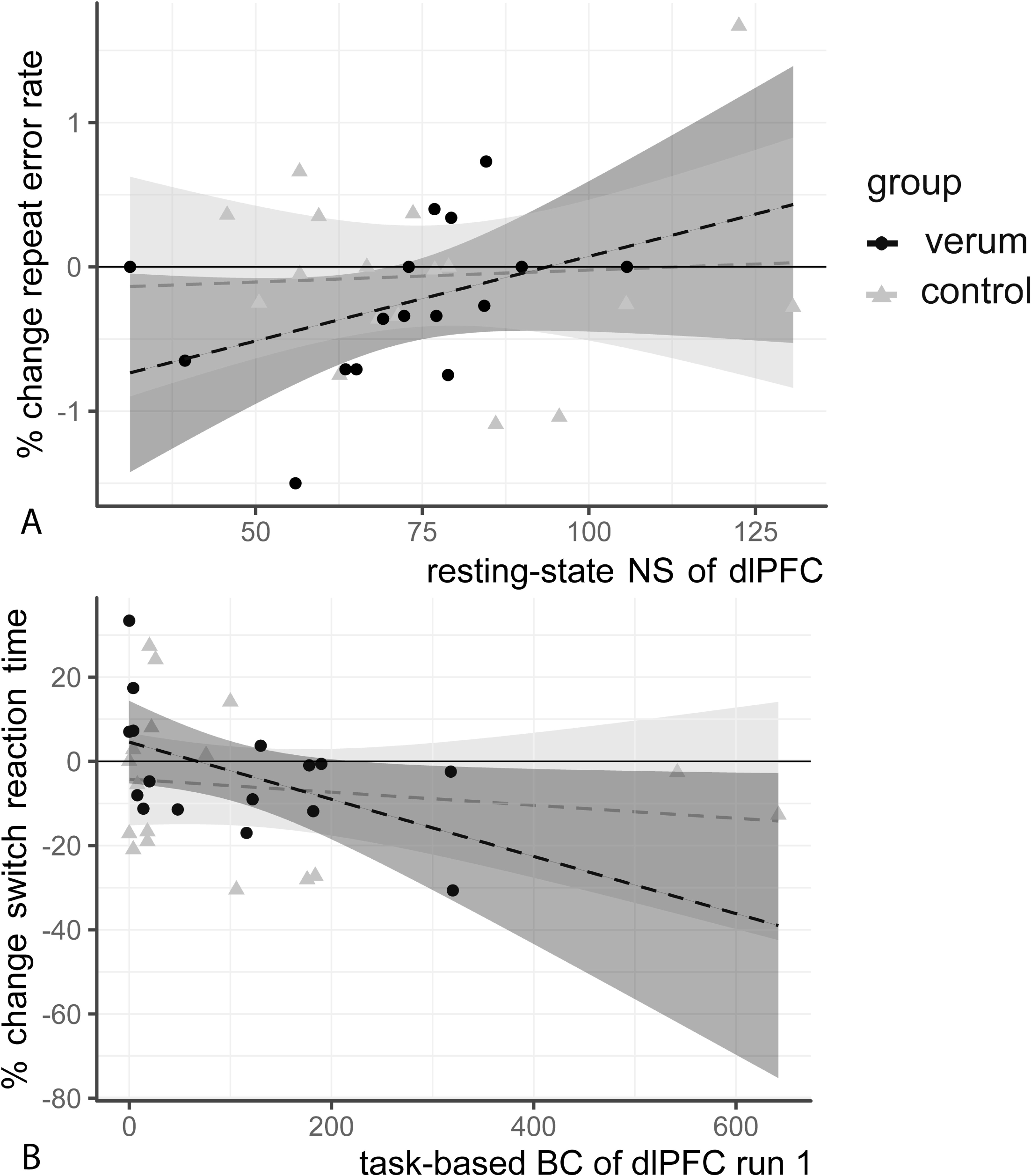
Correlations between baseline fMRI graph measures and change in cognitive performance after LF rTMS: **A.** Higher resting state NS of the dlPFC node is associated with an increase in RER in the verum group (τ= 0.447, p=.017, 95% CI [0.138, 0.781]) but not in the control group (τ= −0.170, p=.343, 95% CI [−0.527, 0.214]; z=2.696, p=0.007). **B.** Lower BC of the dlPFC node is associated with an increase in SRT in the verum group after TMS (τ= −0.403, p= .03, 95% CI [−0.733, −0.031]) but not in the control group (τ = −0.067, p=.710, 95% CI [−0.327, 0.267]; z=−2.040, p=0.04). Shaded areas on plot correspond to (linear) 95% CIs. Note that a linear correlation line and 95% CIs are drawn in these figures, but this association was tested non-parametrically. NS, Node strength; dlPFC, dorsolateral prefrontal cortex; BC, betweenness centrality

### Prediction of behaviour change after rTMS from baseline task-based graph measures

In the verum group, BC of the dlPFC node correlated negatively with percentage change in shift response time (ΔSRT) for both the first (τ= −0.403, p= .030, 95% CI [−0.733, −0.031]) and second run of task-based fMRI (τ=−0.367, p=.048, 95% CI [-.692, 0]) (figure 2b); i.e. the lower the BC, the greater the increase in shift response time after rTMS. This relationship was not seen for BC of the dlPFC in the control group (τ = −0.067, p=.710, 95% CI [−0.327, 0.267]) or for the BC of the control node of the verum group (τ = −0.017, p=.928, 95% CI [−0.482, 0.440]). The correlation in the dlPFC group was also significantly stronger than that of the control group (z=-2.040, p=0.04). Correlations between other task-based graph measures and changes in behaviour were not significant (see Appendix C, table S6).

### Change in task-based graph measures after rTMS

There was no significant difference between verum and control groups in terms of change in network topology after rTMS in either the first or second run of task-based fMRI (see Appendix C, table S7).

### Association of change in behaviour with change in task-based graph measures

In the verum group, percentage change in shift error rate (ΔSER) between sessions was positively correlated with change in BC (ΔBC) of the dlPFC node, but only for the second run (τ = 0.424 p= .024, 95% CI [0.150, 0.669]) (figure 3). In other words, subjects showing an rTMS-induced decrease in BC also had a decreased SER after rTMS. Conversely, those with no change or an increase in BC had an increased SER. There was no significant correlation between ΔSER and ΔBC in the vertex group (τ =−0.180, p=.319, 95% CI: [−0.454, 0.129]) and there was no correlation with ΔBC of the control node (τ = −0.059 p=.752, 95% CI:[−0.581, 0.415]). Furthermore, the correlation coefficients differed significantly between groups (z=2.617, p=0.008). There was no other significant correlation between changes in task based graph measures and changes in behaviour (see Appendix C, table S8).

**Figure 3:**
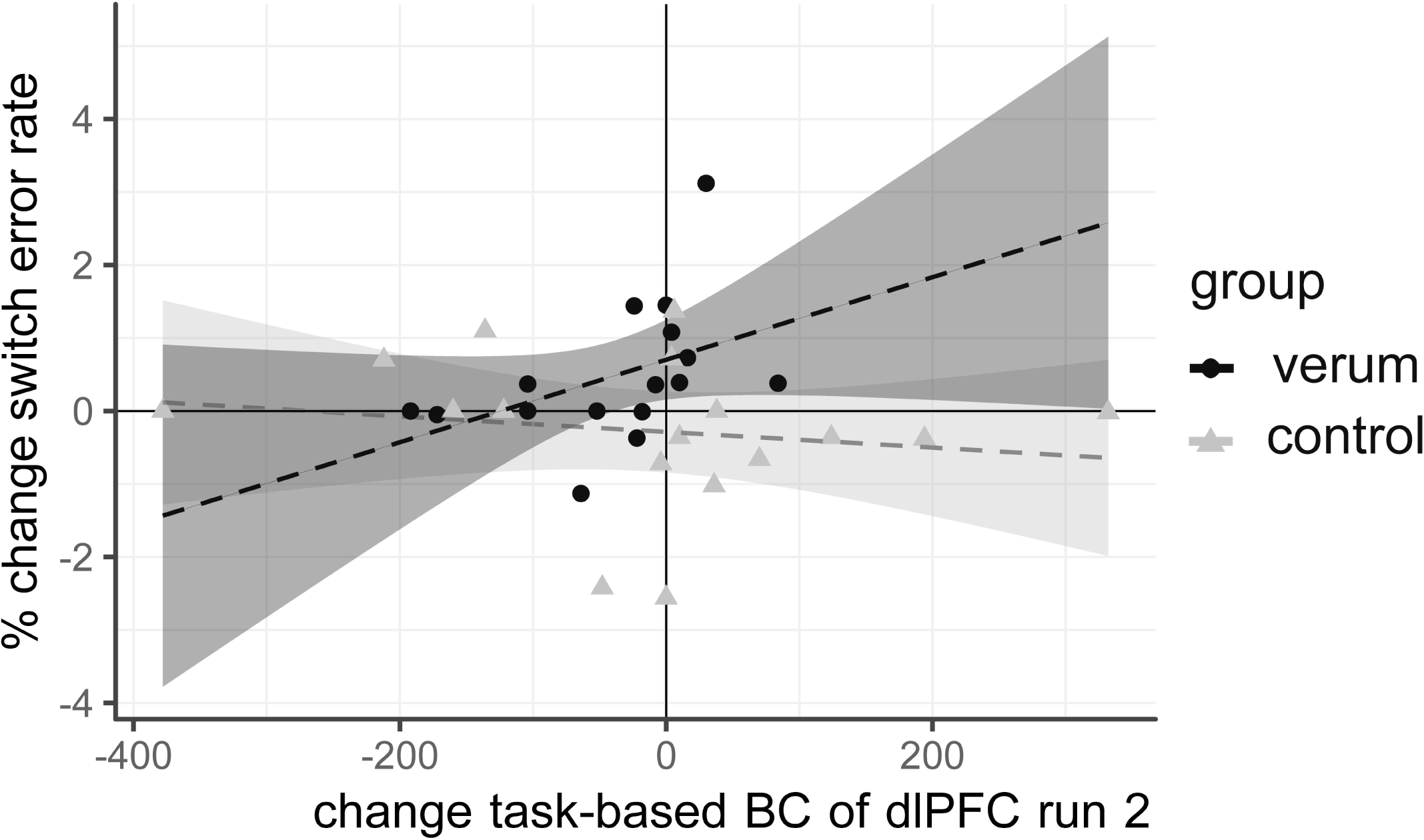
Change in task-based BC of the dLPFC after rTMS is associated with change in shift error rate after rTMS: Participants with no change in task-based BC of the dlPFC after rTMS showed an increase in shift errors, while participants with a decrease in BC after rTMS was showed an improvement or no change in shift errors (verum group: τ = 0.424, p= .024, 95% CI [0.150, 0.669]; control group: τ =−0.180 p=.319, 95% CI:[−0.454, 0.129], z=2.617, p=0.008). Shaded areas on plot correspond to 95% CIs. Note that a linear correlation line and 95% CIs are drawn in these figures, but this association was tested non-parametrically. BC, betweenness centrality; dlPFC, dorsolateral prefrontal cortex.

## Discussion

We investigated whether functional network topology of pre-stimulation rsfMRI or task-based fMRI was predictive of the change in set-shifting performance after inhibitory rTMS to the dlPFC, and whether this change was also related to change in graph features after stimulation. Our findings indicate that individuals with a higher pre-TMS BC of the dlPFC during set-shifting are less affected by inhibitory rTMS, and also show a large decrease in task-based BC following rTMS. Additionally, individuals with a higher pre-TMS NS of the dlPFC during resting state are more affected by inhibitory rTMS.

### Functional network topology and response to LF rTMS

BC gives an indication of the level of global integration of a node. In our study, the detrimental effect of LF rTMS was greater in subjects with a lower pre-TMS BC of the dlPFC, resulting in an rTMS-induced increase in SRT. Conversely, subjects with a higher BC of the dlPFC appeared to be less affected and showed rTMS-induced improvement in SRT. We also found that subjects whose behavioural performance was least affected by the LF rTMS not only had a higher baseline BC of the dlPFC but also showed the greatest rTMS-induced decrease in BC. This suggests that the capacity to ‘lose’ BC and thereby buffer the effects of LF rTMS is a possible determinant of resilience to inhibitory rTMS. In other studies, a high level of network-wide connectivity of the dlPFC has been linked to shorter reaction time in attention tasks[36] and higher fluid intelligence scores[19]. Taken together with our findings, this suggests that a dlPFC that allows more integrated information flow throughout the whole network may provide increased cognitive flexibility and therefore greater capacity to adapt to insults. This apparent resilience of globally well-integrated nodes may only be true in the case of inhibitory rTMS, where brain excitability is temporarily reduced rather than blocked completely: a computational lesion study showed that full deletion of nodes with high BC resulted in global network dysfunction.[17]

We also found an association between higher resting state NS of the dlPFC and rTMS-induced increase in repeat errors. This implies that regions with a higher NS are more vulnerable to the inhibitory effects of rTMS. As NS is a measure of the strength of the connections of a node with its neighbours[12], their loss may lead to network dysfunction[17]. Previous seed-based FC fMRI studies also suggest that the effectiveness of rTMS is dependent on the strength of the connections of the stimulated site; stimulation of regions that have a greater FC at baseline[6,37], results in a greater rTMS response. Inhibitory rTMS may therefore be more effective in subjects with a high baseline resting NS because it allows the effects of inhibition to spread more easily to functionally connected regions.

### Differences between resting state and task based graph measures

Interestingly, different graph measures were associated with a reduction in different aspects of behavioural performance during set-shifting after rTMS in task-based and rsfMRI. BC and ΔBC in task-based scans were specifically associated with change in set shifting performance, while NS in resting state scans was associated with errors on repeat trials. The prominence of BC in the task-based fMRI and NS in rsfMRI could be due to the changes in functional network topology that take place when transitioning from rest to task states. The brain network is more integrated during task execution (with higher BC of the relevant nodes) [38–40],, while measures related to node strength are more important during rest[38,39]. The specificity of task-based scans to set shifting performance rather than repeat trial performance (which may be a measure of more general aspects of cognitive performance such as attention and working memory[41]) is perhaps due to the fact that the stimulation location was chosen from the area of maximum activation on fMRI during set shifting[14]. This indicates that although resting state and task-based networks have a similar intrinsic structure[42], network measures derived from task-based fMRI may have more relevance for predicting rTMS-induced behavioural changes on the same task.

### Limitations and strengths

Our exploratory study has some limitations. Firstly, the relatively small sample size means that our power is limited and any effects present may be inflated[43]. Secondly, the rTMS-induced performance changes are subtle. Tasks with higher cognitive loads or stronger rTMS stimulation may give larger and more robust effects. Thirdly, the interval between pre-rTMS scan and the actual day of rTMS was approximately 2 weeks – it is unknown how representative the networks measured at session 1 were of the pre-stimulation networks actually present at session 2 (though functional connectivity measures have been shown to stay largely stable over time[44,45]). Our cohort also has a wide age range (39-75) and an average age of 56. This may mean that our results are less applicable to younger age groups as graph features have been shown to vary with age[46] (though graph measures showed no correlation in our sample (Appendix C, Table S9)). Replication of this study in a larger cohort with a more demanding task could help tackle these problems. Our study also has a number of strengths. The association of low baseline task-based BC with increase in switch response time was reproduced in the second run of task-based fMRI. The observed effects also survive strict ‘scrubbing’ motion correction.

### Implications and future work

Our results have implications for the selection of rTMS targets. Baseline functional network topology may be an important factor to consider when choosing stimulation targets for therapies involving inhibitory rTMS (for example in the treatment of obsessive-compulsive disorder, stroke, and auditory hallucinations[2]) or in the experimental modelling of the cognitive deficits seen in these diseases in healthy controls. Future work could explore whether the effects demonstrated in this exploratory study are consistent across other stimulation sites, for other cognitive tasks, or for clinical improvement in disease states; which graph measures predict the outcome of excitatory rTMS; and whether specifically targeting nodes with high NS or low BC results in more consistent rTMS outcome.

## Conclusions

We have shown that changes in cognitive performance after inhibitory rTMS to the dlPFC are associated with baseline resting state and task-based functional network graph measures. Subjects with stimulated regions that are globally well-connected during a task are more resilient to the effects of inhibitory rTMS, while those with stimulated regions with strong local connections in the resting state are more vulnerable to inhibitory rTMS. These results have important implications for our understanding of individual variability in response to rTMS, and for the practical application of this non-invasive brain stimulation technique.

## Acknowledgements

We would like to thank Dr. Niels Gerrits for his help with data collection and prof. dr. Jos Twisk for advice on statistical analysis.

## Abbreviations

BC: betweenness centrality
dlPFC: dorsolateral prefrontal cortex
FC: functional connectivity
HF: high frequency
LF: low frequency
MMSE: Mini Mental State Examination
NS: node strength
PC: participation coefficient
RER: repeat error rate
RRT: repeat reaction time
rTMS: repetitive transcranial magnetic stimulation
SER: shift error rate
SRT: shift reaction time

