## Appendix for "Resting-state and task-based centrality of dorsolateral prefrontal cortex predict resilience to inhibitory repetitive transcranial magnetic stimulation"

**Appendix A: Image preprocessing steps**

All image processing was performed using FMRIB's software library (FSL) version 5.0.8

- Removal of non-brain tissue using the Brain Extraction Tool[1],
- Grey and white matter segmentation using FAST[2].
- Parcellation into 225 regions in order to define nodes for graph analysis. We used the Brainnetome Atlas[3] to define 210 cortical regions; 14 subcortical areas were individually segmented using FSL FIRST[4]; and one cerebellar ROI from FSL's cerebellar atlas was segmented[5].
- Preprocessing of fMRI data using the FSL MELODIC pipeline[6]: discarding the first four volumes, motion correction, spatial smoothing (5 mm full width at half maximum), and high-pass filtering (100 second cut-off).
- Coregistration to T1-weighted anatomical MR scans using linear and non-linear co-registration methods[7].
- To account for EPI distortions near air/tissue boundaries during scanning, we applied a mask to the functional scan to exclude voxels with signal intensities in the lowest quartile of the robust range[8]. Any nodes containing <4 voxels were excluded. A total of 193 regions common to all fMRI runs remained: excluded regions were located in the orbitofrontal gyrus, inferior temporal gyrus, parahippocampal gyrus, and thalamus.
- Extraction of time-series from each parcellated region.

### Appendix B: Subdivision into dlPFC-containing modules

Networks can be subdivided into groups of regions that are densely interconnected, known as modules or subnetworks. In the context of functional networks, they are thought to represent areas of segregated processing[9], and specific modules may correspond to specific cognitive functions[10]. The Louvain Modularity algorithm from the Brain Connectivity Toolbox[9] (https://sites.google.com/site/bctnet/) was used to define dlPFC-containing modules for each participant individually, and for RS and each task-based run separately, as modularity is known to vary between individuals and between task and resting states[11]. Since Louvain Modularity is an heuristic algorithm, and produces slightly different subdivisions of nodes each time it is run[12], we permuted the algorithm 1000 times, calculated pairwise similarities between all iterations, and used the iteration with the most similarity to all other iterations as a consensus network[13]. A resolution parameter (gamma) of 1.1 was used as this gave on average 7 modules per participant, which is a previously described stable subdivision of known resting state functional networks[14].

### Appendix C: Supplementary data tables

#### Table S1: Group comparisons of set shifting performance at baseline

| | Verum | Control | *p* |
|---|---|---|---|
| SER (%) | 0.41 ± 0.61 | 0.68 ± 1.01 | .444[a] |
| RER (%) | 1.12 ± 1.21 | 1.03 ± 1.11 | .873[a] |
| SRT (s) | 927.40 ± 230.46 | 901.40 ± 210.00 | .737[b] |
| RRT (s) | 849.50 ± 211.60 | 819.70 ± 201.30 | .681[b] |

[a]Mann-Whitney U test; [b]Independent samples t-test. Data presented as mean ± SD. Abbreviations: SER: switch error rate; RER: repeat error rate; SRT: switch reaction time; RRT: repeat reaction time.

#### Table S2: Correlations between potential confounders and change in set shifting performance after rTMS

| | IQ | | | Educational level | | | MMSE | | | rTMS-task interval | | |
|---|---|---|---|---|---|---|---|---|---|---|---|---|
| | Tau | 95% CI | | Tau | 95% CI | | Tau | 95% CI | | Tau | 95% CI | |
| | | LB | UB | | LB | UB | | LB | UB | | LB | UB |
| ∆SER | -0.067 | -0.357 | 0.235 | 0.141 | -0.170 | 0.464 | 0.179 | -0.118 | 0.435 | 0.019 | -0.250 | 0.275 |
| ∆RER | -0.006 | -0.293 | 0.270 | -0.030 | -0.298 | 0.230 | 0.141 | -0.143 | 0.430 | 0.059 | -0.251 | 0.339 |
| ∆RRT | -0.027 | -0.285 | 0.245 | -0.215 | -0.444 | 0.041 | -0.212 | -0.452 | 0.051 | 0.142 | -0.173 | 0.446 |
| ∆SRT | -0.112 | -0.356 | 0.137 | -0.047 | -0.319 | 0.241 | -0.062 | -0.270 | 0.141 | 0.157 | -0.140 | 0.473 |

Kendall's Tau-b correlations with bootstrapped confidence intervals. Abbreviations: ∆SER: change in switch error rate; ∆RER: change in repeat error rate; ∆SRT: change in switch reaction time; ∆RRT: change in repeat reaction time; CI: confidence interval; LB: lower bound; UB: upper bound; MMSE = Mini-Mental State Examination.

#### Table S3: rsfMRI baseline graph measures – comparison between groups

| Graph measure | Verum | | Control | | *p* |
|---|---|---|---|---|---|
| PC of dlPFC | 0.667 | ± 0.068 | 0.640 | ± 0.046 | .146 |
| BC of dlPFC | 106.875 | ±124.915 | 71.058 | ± 80.960 | .683 |
| NS of dlPFC | 71.634 | ±18.317 | 76.923 | ± 24.302 | 1.000 |

Data presented as mean ± SD. Group comparisons carried out using Mann-Whitney U tests. Abbreviations: dlPFC: dorsolateral prefrontal cortex; BC: betweenness centrality; PC: participation coefficient; NS: node strength.

#### Table S4: Correlations between all baseline resting state fMRI graph measures and change in all behavioural outcomes for dlPFC and vertex groups

| | ∆SER | | | ∆RER | | | ∆RRT | | | ∆SRT | | |
|---|---|---|---|---|---|---|---|---|---|---|---|---|
| | Tau | 95% CI | | Tau | 95% CI | | Tau | 95% CI | | Tau | 95% CI | |
| | | LB | UB | | LB | UB | | LB | UB | | LB | UB |
| Verum group | | | | | | | | | | | | |
| PC of dlPFC | 0.076 | -0.346 | 0.457 | -0.329 | -0.715 | 0.066 | -0.050 | -0.413 | 0.279 | 0.050 | -0.407 | 0.440 |
| BC of dlPFC | -0.119 | -0.474 | 0.284 | -0.288 | -0.596 | 0.070 | -0.293 | -0.689 | 0.170 | -0.259 | -0.621 | 0.200 |
| NS of dlPFC | -0.076 | -0.547 | 0.452 | 0.447 | 0.138 | 0.781 | 0.217 | -0.232 | 0.637 | 0.117 | -0.342 | 0.571 |
| Control group | | | | | | | | | | | | |
| PC of dlPFC | -0.045 | -0.425 | 0.335 | 0.288 | -0.052 | 0.570 | .368 | 0.027 | 0.694 | .471 | 0.163 | 0.767 |
| BC of dlPFC | -0.060 | -0.598 | 0.458 | 0.244 | -0.121 | 0.578 | 0.191 | -0.143 | 0.446 | 0.088 | -0.279 | 0.396 |
| NS of dlPFC | -0.196 | -0.597 | 0.228 | -0.170 | -0.527 | 0.214 | -0.044 | -0.341 | 0.254 | -0.147 | -0.541 | 0.253 |

Kendall's Tau-b correlations with bootstrapped confidence intervals. Abbreviations: dlPFC: dorsolateral prefrontal cortex; BC: betweenness centrality; PC: participation coefficient; NS: node strength. ∆SER: change in switch error rate; ∆RER: change in repeat error rate; ∆SRT: change in switch reaction time; ∆RRT: change in repeat reaction time; CI: confidence interval; LB: lower bound; UB: upper bound,

#### Table S5: Baseline task-based graph measures – comparison between groups

#### a. Run 1

| Graph measure | Verum | | Control | | *p* |
|---|---|---|---|---|---|
| PC of dlPFC | 0.671 | ± 0.059 | 0.617 | ± 0.074 | .031 |
| BC of dlPFC | 103.375 | ±110.056 | 114.47<br>0 | ± 189.953 | .709 |
| NS of dlPFC | 81.303 | ±18.529 | 84.572 | ± 23.812 | .657 |

Data presented as mean ± SD. Group comparisons carried out using Mann-Whitney U tests. Abbreviations: dlPFC: dorsolateral prefrontal cortex; BC: betweenness centrality; PC: participation coefficient; NS: node strength

#### b. Run 2

| Graph measure | Verum | | Control | | *p* |
|---|---|---|---|---|---|
| PC of dlPFC | 0.659 | ± 0.066 | 0.655 | ± 0.088 | .986 |
| BC of dlPFC | 81.500 | ± 80.031 | 139.52<br>9 | ± 206.915 | .901 |
| NS of dlPFC | 78.841 | ±23.933 | 80.958 | ± 24.912 | .845 |

Data presented as mean ± SD. Group comparisons carried out using Mann-Whitney U tests. Abbreviations: dlPFC: dorsolateral prefrontal cortex; BC: betweenness centrality; PC: participation coefficient; NS: node strength

### Table S6: Correlations between all baseline task-based fMRI graph measures and change in all behavioural outcomes for dlPFC and vertex groups

#### c. Run 1

| | ΔSER | | | ΔRER | | | ΔRRT | | | ΔSRT | | |
|---|---|---|---|---|---|---|---|---|---|---|---|---|
| | Tau | 95% CI | | Tau | 95% CI | | Tau | 95% CI | | Tau | 95% CI | |
| | | LB | UB | | LB | UB | | LB | UB | | LB | UB |
| Verum group | | | | | | | | | | | | |
| PC of dlPFC | 0.000 | -0.416 | 0.448 | -0.119 | -0.485 | 0.302 | -0.042 | -0.453 | 0.396 | -0.142 | -0.559 | 0.336 |
| PC control node | 0.000 | -0.465 | 0.492 | -0.254 | -0.697 | 0.132 | -0.025 | -0.501 | 0.488 | -0.059 | -0.445 | 0.374 |
| BC of dlPFC | 0.093 | -0.380 | 0.523 | 0.008 | -0.295 | 0.343 | -0.217 | -0.544 | 0.123 | -0.117 | -0.424 | 0.210 |
| BC control node | -0.127 | -0.560 | 0.310 | 0.228 | -0.094 | 0.575 | 0.183 | -0.172 | 0.546 | 0.050 | -0.394 | 0.492 |
| NS of dlPFC | -0.228 | -0.566 | 0.160 | -0.008 | -0.337 | 0.316 | 0.250 | -0.125 | 0.629 | 0.117 | -0.334 | 0.497 |
| NS control node | 0.009 | -0.434 | 0.458 | 0.213 | -0.183 | 0.573 | 0.286 | -0.146 | 0.699 | 0.168 | -0.357 | 0.592 |
| Control group | | | | | | | | | | | | |

| | | | | | | | | | | | | |
|---|---|---|---|---|---|---|---|---|---|---|---|---|
| PC of dlPFC | -0.068 | -0.509 | 0.372 | -0.059 | -0.369 | 0.267 | 0.081 | -0.342 | 0.437 | -0.140 | -0.559 | 0.236 |
| BC of dlPFC | 0.236 | -0.072 | 0.527 | 0.015 | -0.353 | 0.386 | -0.037 | -0.411 | 0.361 | -0.067 | -0.327 | 0.267 |
| BC control node | 0.165 | -0.350 | 0.625 | 0.066 | -0.356 | 0.476 | 0.074 | -0.333 | 0.485 | 0.000 | -0.493 | 0.491 |
| NS of dlPFC | -0.211 | -0.609 | 0.248 | -0.052 | -0.506 | 0.378 | 0.088 | -0.319 | 0.461 | 0.103 | -0.384 | 0.518 |
| NS control node | -0.190 | -0.620 | 0.266 | -0.149 | -0.507 | 0.224 | 0.022 | -0.386 | 0.387 | 0.082 | -0.425 | 0.505 |

Kendall's Tau-b correlations with bootstrapped confidence intervals. Abbreviations: dlPFC: dorsolateral prefrontal cortex; BC: betweenness centrality; PC: participation coefficient; NS: node strength. ΔSER: change in switch error rate; ΔRER: change in repeat error rate; ΔSRT: change in switch reaction time; ΔRRT: change in repeat reaction time; CI: confidence interval; LB: lower bound; UB: upper bound

#### d. Run 2

| | ΔSER | | | ΔRER | | | ΔRRT | | | ΔSRT | | |
|---|---|---|---|---|---|---|---|---|---|---|---|---|
| | Tau | 95% CI | | Tau | 95% CI | | Tau | 95% CI | | Tau | 95% CI | |
| | | LB | UB | | LB | UB | | LB | UB | | LB | UB |
| Verum group | | | | | | | | | | | | |
| PC of dlPFC | -0.177 | -0.622 | 0.368 | -0.228 | -0.706 | 0.301 | -0.017 | -0.379 | 0.325 | -0.017 | -0.377 | 0.326 |
| PC control node | -0.110 | -0.540 | 0.387 | -0.228 | -0.667 | 0.241 | -0.017 | -0.396 | 0.344 | -0.017 | -0.323 | 0.306 |
| BC of dlPFC | -0.143 | -0.653 | 0.322 | -0.228 | -0.509 | 0.085 | -0.183 | -0.511 | 0.168 | -0.150 | -0.534 | 0.286 |
| BC control node | 0.110 | -0.292 | 0.605 | 0.228 | -0.105 | 0.529 | 0.183 | -0.197 | 0.569 | 0.250 | -0.169 | 0.656 |
| NS of dlPFC | 0.111 | -0.257 | 0.498 | 0.145 | -0.217 | 0.463 | 0.151 | -0.182 | 0.513 | 0.151 | -0.246 | 0.536 |
| NS control node | 0.177 | -0.245 | 0.654 | 0.228 | -0.143 | 0.550 | 0.150 | -0.280 | 0.564 | 0.150 | -0.333 | 0.580 |
| Control group | | | | | | | | | | | | |
| PC of dlPFC | 0.000 | -0.454 | 0.423 | 0.288 | -0.062 | 0.605 | -0.176 | -0.554 | 0.279 | 0.015 | -0.367 | 0.398 |
| BC of dlPFC | 0.197 | -0.283 | 0.633 | 0.201 | -0.193 | 0.579 | -0.030 | -0.419 | 0.388 | -0.074 | -0.452 | 0.342 |
| BC control node | 0.135 | -0.337 | 0.557 | 0.066 | -0.331 | 0.444 | 0.132 | -0.245 | 0.576 | 0.059 | -0.365 | 0.493 |
| NS of dlPFC | -0.211 | -0.601 | 0.225 | 0.022 | -0.335 | 0.412 | -0.044 | -0.492 | 0.340 | 0.029 | -0.455 | 0.458 |
| NS control node | -0.143 | -0.588 | 0.335 | -0.074 | -0.437 | 0.355 | -0.111 | -0.474 | 0.182 | -0.037 | -0.467 | 0.369 |

Kendall's Tau-b correlations with bootstrapped confidence intervals. Abbreviations: dlPFC: dorsolateral prefrontal cortex; BC: betweenness centrality; PC: participation coefficient; NS: node strength. ΔSER: change in switch error rate; ΔRER: change in repeat error rate; ΔSRT: change in switch reaction time; ΔRRT: change in repeat reaction time; CI: confidence interval; LB: lower bound; UB: upper bound

### Table S7: Mean % change from baseline of task based graph measures after rTMS: comparison between groups

| | Run 1 | | | Run 2 | | |
|---|---|---|---|---|---|---|
| | Verum | Control | *p* | Verum | Control | *p* |
| PC of dlPFC | -3.493 ± 11.863 | 2.368 ± 15.049 | .191 | -2.952 ± 13.120 | -3.44 ± 12.769 | .533 |
| BC of dlPFC [a] | -16.267 ± 130.933 | 4.118 ± 151.249 | .326 | -38.500 ± 73.457 | -14.471 ± 161.361 | .363 |
| NS of dlPFC | -8.556 ± 23.921 | 6.052 ± 22.061 | .087 | 5.755 ± 40.998 | 15.892 ± 26.026 | .168 |

Data presented as mean % change from baseline ± SD. Group comparisons carried out using Mann-Whitney U tests. [a]Presented as absolute change from baseline to avoid dividing by zero. Abbreviations: dlPFC: dorsolateral prefrontal cortex; BC: betweenness centrality; PC: participation coefficient; NS: node strength

### Table S8: correlations between change in all task based fMRI graph measures and change in all behavioural outcomes for dlPFC and vertex groups

#### a. Run 1

| | ΔSER | | | ΔRER | | | ΔRRT | | | ΔSRT | | |
|---|---|---|---|---|---|---|---|---|---|---|---|---|
| | Tau | 95% CI | | Tau | 95% CI | | Tau | 95% CI | | Tau | 95% CI | |
| | | LB | UB | | LB | UB | | LB | UB | | LB | UB |
| Verum group | | | | | | | | | | | | |
| ΔPC of dlPFC | -0.262 | -0.596 | 0.100 | 0.059 | -0.334 | 0.481 | 0.033 | -0.346 | 0.445 | 0.033 | -0.372 | 0.394 |
| ΔBC of dlPFC | 0.211 | -0.247 | 0.606 | 0.262 | -0.116 | 0.617 | 0.233 | -0.154 | 0.552 | 0.033 | -0.472 | 0.477 |
| ΔNS of dlPFC | 0.110 | -0.323 | 0.598 | 0.093 | -0.209 | 0.419 | -0.033 | -0.425 | 0.333 | 0.033 | -0.353 | 0.494 |
| Control group | | | | | | | | | | | | |
| ΔPC of dlPFC | 0.120 | -0.277 | 0.513 | 0.125 | -0.230 | 0.427 | 0.029 | -0.333 | 0.411 | 0.309 | -0.119 | 0.688 |
| ΔBC of dlPFC | -0.219 | -0.520 | 0.155 | 0.252 | -0.131 | 0.687 | -0.096 | -0.459 | 0.240 | -0.155 | -0.480 | 0.187 |
| ΔNS of dlPFC | 0.090 | -0.332 | 0.457 | 0.170 | -0.261 | 0.541 | 0.015 | -0.371 | 0.388 | 0.059 | -0.396 | 0.482 |

Kendall's Tau-b correlations with bootstrapped confidence intervals. Abbreviations: dlPFC: dorsolateral prefrontal cortex; ΔBC: change in betweenness centrality; ΔPC: change in participation coefficient; NS: change in node strength. ΔSER: change in switch error rate; ΔRER: change in repeat error rate; ΔSRT: change in switch reaction time; ΔRRT: change in repeat reaction time; CI: confidence interval; LB: lower bound; UB: upper bound

#### b. Run 2

| | ΔSER | | | ΔRER | | | ΔRRT | | | ΔSRT | | |
|---|---|---|---|---|---|---|---|---|---|---|---|---|
| | Tau | 95% CI | | Tau | 95% CI | | Tau | 95% CI | | Tau | 95% CI | |
| | | LB | UB | | LB | UB | | LB | UB | | LB | UB |
| DlPFC group | | | | | | | | | | | | |
| ΔPC of dlPFC | 0.042 | -0.367 | 0.450 | 0.110 | -0.430 | 0.633 | -0.167 | -0.531 | 0.194 | -0.233 | -0.524 | 0.087 |
| ΔBC of dlPFC | 0.424 | 0.150 | 0.699 | 0.288 | -0.181 | 0.688 | 0.377 | -0.046 | 0.740 | 0.243 | -0.140 | 0.651 |
| ΔNS of dlPFC | -0.059 | -0.515 | 0.390 | 0.025 | -0.400 | 0.431 | -0.067 | -0.399 | 0.265 | -0.200 | -0.552 | 0.120 |
| Vertex group | | | | | | | | | | | | |
| ΔPC of dlPFC | 0.105 | -0.312 | 0.505 | -0.185 | -0.581 | 0.215 | 0.074 | -0.260 | 0.537 | 0.059 | -0.333 | 0.466 |
| ΔBC of dlPFC | -0.180 | -0.454 | 0.129 | 0.170 | -0.266 | 0.542 | 0.015 | -0.411 | 0.440 | 0.000 | -0.422 | 0.395 |
| ΔNS of dlPFC | 0.196 | -0.137 | 0.503 | -0.111 | -0.559 | 0.365 | -0.029 | -0.463 | 0.379 | -0.015 | -0.369 | 0.346 |

Kendall's Tau-b correlations with bootstrapped confidence intervals. Abbreviations: dlPFC: dorsolateral prefrontal cortex; ΔBC: change in betweenness centrality; ΔPC: change in participation coefficient; NS: change in node strength. ΔSER: change in switch error rate; ΔRER: change in repeat error rate; ΔSRT: change in switch reaction time; ΔRRT: change in repeat reaction time; CI: confidence interval; LB: lower bound; UB: upper bound

### Table S9: Correlations between age and resting state graph measures at baseline

| | Age | | |
|---|---|---|---|
| | Tau | 95% CI | |
| | | LB | UB |
| PC of dlPFC | 0.006 | -0.232 | 0.248 |
| BC of dlPFC | 0.108 | -0.160 | 0.370 |
| NS of dlPFC | -0.021 | -0.287 | 0.269 |

Kendall's Tau-b correlations with bootstrapped confidence intervals. Abbreviations: dlPFC: dorsolateral prefrontal cortex; BC: betweenness centrality; PC: participation coefficient; NS: node strength
